# The bidirectional role of GABA_A_ and GABA_B_ receptors during the differentiation process of neural precursor cells of the subventricular zone

**DOI:** 10.1101/2024.03.19.585835

**Authors:** Estefanía Gutierrez-Castañeda, Vladimir Alex Martínez-Rojas, Lenin David Ochoa-de la Paz, Emilio J. Galván

## Abstract

The intricate process of neuronal differentiation integrates multiple signals to induce transcriptional, morphological, and electrophysiological changes that reshape the properties of neural precursor cells during their maturation and migration process. An increasing number of neurotransmitters and biomolecules have been identified that serve as molecular signals that trigger and guide this process. In this sense, taurine, a sulfur-containing, non-essential amino acid widely expressed in the mammal brain, modulates the neuronal differentiation process. In this study, we describe the effect of taurine acting via the ionotropic GABA_A_ receptor and the metabotropic GABA_B_ receptor on the neuronal differentiation and electrophysiological properties of precursor cells derived from the subventricular zone of the mouse brain. Taurine stimulates the number of neurites and favors the dendritic complexity of the neural precursor cells, accompanied by changes in the somatic input resistance and the strength of inward and outward membranal currents. At the pharmacological level, the blockade of GABA_A_ receptors inhibits these effects, whereas the stimulation of GABA_B_ receptors has no positive effects on the taurine-mediated differentiation process. Strikingly, the blockade of the GABA_B_ receptor with CGP533737 stimulates neurite outgrowth, dendritic complexity, and membranal current kinetics of neural precursor cells. The effects of taurine on the differentiation process involve Ca^2+^ mobilization and the activation of intracellular signaling cascades since chelation of intracellular calcium with BAPTA-AM, and inhibition of the CaMKII, ERK1/2, and Src kinase inhibits the neurite outgrowth of neural precursor cells of the subventricular zone.

## Introduction

Adult neurogenesis, the process by which newborn neurons integrate into consolidated neural circuits, has been extensively studied in the subgranular zone of the dentate gyrus and the subventricular zone (SVZ) of the lateral ventricles. Neurons stemming from the subgranular zone merge into the granule cell layer and develop a glutamatergic phenotype, contributing to hippocampal-related functions (Akers et al., 2014; Altman & Das, 1965; Snyder et al., 2011; Zhao et al., 2006). On the other hand, most of the SVZ-derived neurons undergo a rostral migration that culminates with their integration into the olfactory bulb. These newborn cells develop a GABAergic phenotype and are, indeed, interneurons (Inta et al., 2008; Lois & Alvarez-Buylla, 1994) whose function has been linked to accelerating the learning of olfactory discrimination and short-term olfactory memory (Alonso et al., 2012; Sahay et al., 2011).

Morphologically, adult neurogenesis is accompanied by neurite outgrowth and the formation of a multifarious neurite branching rearrangement associated with neuronal maturation (Zhang et al., 2016). Neurite outgrowth is triggered by depolarizing events that favor transient increases in intracellular calcium and the activation of signaling cascades that promote neurite elongation and synapse formation (Sogawa et al., 2000). Consistent with this tenet, neuronal precursor cells (NPCs) of the SVZ are stem cells endowed with multiple signaling cascades whose activation converges in the phosphorylation of the cAMP response element-binding protein (CREB), a transcription factor required for neurite outgrowth of NPCs (Gampe et al., 2011; Herold et al., 2011).

The signaling cascades involved in the differentiation process of NPCs are activated by diverse neurotransmitters, hormones, and biomolecules, including taurine, a free sulfur-amino acid found in high concentrations in the mammalian brain (Huxtable, 1992). It has been well established that taurine has a neurotrophic effect on neurite outgrowth and the thickness of developing neurites (Mersman et al., 2020). In a recent study, we demonstrated that the development of neurite complexity of NPC SVZ mediated by taurine is accompanied by the functional expression of ion channels that reshape the passive electrophysiological properties of NPC SVZ and favor the appearance of active properties, including action potential generation and a Ca^2+^-dependent afterhyperpolarization (Gutierrez-Castañeda et al.).

Interestingly, a series of studies have documented that NPCs also express GABA_B_ receptors (GABA_B_R). The blockade of GABA_B_R promotes hippocampal neurogenesis (Felice et al., 2012), and in neocortical-derived neurospheres, the blockade of GABA_B_R favors the proliferation and differentiation process (Fukui et al., 2008). Contrary to these findings, the pharmacological activation of GABA_B_R has been reported to suppress adult hippocampal neurogenesis (Giachino et al., 2014; Gustorff et al., 2021). Despite the possible relevance of GABA_B_R in the differentiation process of NPCs, the expression of GABA_B_R on NPC SVZ and its role in the differentiation process of NPC SVZ remains unexplored.

To close this gap, in this study, we explored the interaction of taurine with GABA receptors in the maturation process of NPC SVZ. We first demonstrated the presence of GABA_B_R in neurospheres of NPC SVZ; then, we explored the role of taurine in the establishment of inward and outward membrane currents that shape the electrophysiological properties associated with neuronal maturation; and finally, we demonstrated the role of several intracellular signaling cascades in the neurite formation of NPC SVZ. Our findings demonstrate that the abundant neuroactive molecule taurine plays a significant role in the neurophysiological maturation of NPC SVZ via an antagonistic modulation of ionotropic and metabotropic GABA receptors acting in a nonsynaptic fashion.

## Methods

### Primary neural precursor cell culture

The experimental procedures were conducted in strict accordance with the Mexican Official Norm for the use and care of laboratory animals “NOM-062-ZOO-1999,” the Guide for the Care and Use of Laboratory Animals of the National Institutes of Health, and with the approval of the Institutional Animal Care and Use Committee of the National University of Mexico (FMED/DI/068/2019, CICUAL018-CIC-2019). Eight-day-old CD1 mice were used to obtain NPC SVZ. The mice were sacrificed by decapitation, their brain was removed, and the SVZ was isolated. Then, the tissue was mechanically dissociated and placed in a DMEM/F12 medium (Dulbecco’s Modified Eagle Medium/Nutrient Mixture F-12. Gibco; Thermo Fisher Scientific, Inc., Waltham, MA, USA) and centrifuged for 5 min at 1000 rpm. The supernatant was removed, and the pellet was resuspended and cultured in a growth medium (DMEM/F12 supplemented with B27, plus epidermal growth factor and fibroblast growth factor 2) to promote progenitor cell proliferation and SVZ neurosphere formation.

The resulting cultures were maintained at 37°C and 5% CO_2_. After seven days, the primary neurospheres were mechanically disaggregated with trypsin, and a trypsin inhibitor (1:1) was added to stop the reaction. The cells were then cultured in a growth medium for four to five days at 37°C and 5% CO_2_. For the differentiation process of NPC SVZ, disaggregated cells of the secondary SVZ neurospheres were cultured in 48 wells pretreated with poly-D-lysine (Sigma-Aldrich, St. Louis, MO, USA) and a differentiation medium (DMEM/F12 supplemented with 1% fetal bovine serum (FBS) and complemented with taurine (10 mM)). The taurine dose used in this study was previously determined by performing a dose-response curve (see Gutierrez-Castañeda et al., 2023).

To evaluate the role of the GABA_A_ or GABA_B_ receptor in the differentiation process, the cultured cells were preincubated for one hour with the GABA_A_ receptor antagonist picrotoxin (100 µM), the GABA_B_ receptor antagonist CGP55845 (5 µM), or the GABA_B_ receptor agonist baclofen (100 µM). To determine the participation of different signaling cascades in the differentiation process induced by taurine through GABA receptors, the cells were cultured for differentiation for three days and preincubated with BAPTA-AM (100 µM) in a Krebs-Ringer Buffer without calcium for 30 min. Then, KN93 (10 µM), FR180204 (10 µM), and SrcI (1 µM) were co-incubated in the presence of taurine 10 (mM) for seven days. All the drugs were purchased from Sigma-Aldrich, St. Louis, MO, USA.

### Immunofluorescence assay

The cultured cells were fixed with cold paraformaldehyde (4%), washed with phosphate-buffered saline (PBS) + 0.1% bovine serum albumin (BSA; 3 × 5 min), and permeabilized/blocked with PBS + 0.1% BSA + 10% FBS + 0.3% Triton X-100 for 1 h at room temperature. The cells were subsequently incubated overnight with the primary antibodies nestin (Cell Signaling Technology Cat# 4760, RRID: AB_2235913) [1:300]), KI67 (Cell Signaling Technology Cat# 9027, RRID:AB_2636984) [1:100), GABABR1(Santa Cruz Biotechnology Cat# sc-166408, RRID:AB_2108175), doublecortin (DCX, Abcam Cat# ab18723, RRID: AB_732011) [1:1000]), and MAP2 (Cell Signaling Technology Cat# 8707, RRID: AB_2722660). The primary antibodies were removed the following day, followed by a 1-h incubation with the secondary antibodies Alexa Fluor-488 anti-rabbit 9 and Alexa Fluor-647 anti-mouse. The excess secondary antibody was then washed and removed from the preparation assembly. The nuclei were stained with DAPI (20 µg/mL) included in the mounting solution (100/500 µL).

### Quantitative analysis for cell markers

Microphotographs were obtained with the cell imaging multi-mode plate reader Cytation™ 5 (BioTek Instruments Inc., Winooski, VT, USA) using a green fluorescent protein (GFP) filter cube (excitation 469/35 nm; emission 525/39 nm; dichroic mirror 497 nm) and 465 nm LED; a Texas Red filter cube (excitation 586/15 nm; emission 647/67 nm; dichroic mirror 605 nm) and 590 nm LED; and a 4′,6-diamidino-2-phenylindole (DAPI) filter cube (excitation 377/50 nm; emission 447/60 nm; dichroic mirror 409) and 365 nm LED. Microphotographs of five fields were randomly selected from each coverslip, and the number of total (DAPI positive) cells and cells positive for the different markers in the selected fields were quantified. Cell images were processed using Gen5™ software (BioTek Instruments Inc.). The results were expressed as the percentage of positive cells out of the total number of cells.

### Morphometric analysis of dendritic arborization of DCX- and MAP2-positive cells

Microphotographs of DCX-positive (DCX+) cells were randomly obtained with a 20x magnification lens. DCX+ cells were analyzed for each experimental condition. The number of primary, secondary, and tertiary neurites and the length of the primary neurites were determined with Gen 5 3.04. The difference between primary, secondary, and tertiary neurites was analyzed in the different experimental conditions. Manual quantification of dendritic arborization was performed by counting the number of primary dendrites for each neuron using ImageJ software (Schneider et al., 2012) with the Sholl analysis plug-in v3.4.2 (Ferreira et al., 2014). Primary neurites were defined as those that originated directly from the soma; secondary and tertiary neurites were defined as those that originated from primary and secondary neurites, respectively. Researchers were blinded to experimental groups prior to performing all morphometric analyses.

### Whole-cell patch-clamp recordings

Dissociated NPC SVZ from SVZ neurospheres were taken from a petri dish containing the cell culture, mounted on a coverslip, and transferred to a submersion recording chamber. They were visualized with infrared differential interference contrast optics coupled to a Nikon FN1 microscope (Nikon Corporation, Minato, Tokyo, Japan). The NPC-SVZ were continuously perfused with Ringer’s solution at the rate of 3–4 mL/min^-1^ with the help of a peristaltic pump (120S, Watson-Marlow, Wilmington, MA, USA). The composition of the bath solution was as follows (mM): 132 NaCl, 3.6 KCl, 1.5 MgCl_2_, 2.5 CaCl_2_, 10 HEPES, and 10 D-glucose; pH = 7.30– 7.40; osmolarity = 280–290 mOsm. The recording chamber temperature was maintained at 32 ± 1°C with a single channel temperature controller (TC-324C, Warner Instruments, Hamden, CT, USA). The patch pipettes were pulled from borosilicate glasses using a micropipette puller (P97, Sutter Instruments, Novato, CA, USA). The puller was programmed to obtain pipettes with a tip resistance of 3– 5 MΩ when filled with an intracellular solution containing the following (mM): 140 KCl, 1.1 EGTA, 10 HEPES, 3 Mg^2+^-ATP, 0.3 Na^+^-GTP, and pH = 7.20–7.30. Whole-cell recordings were performed with an Axopatch 200B amplifier (Molecular Devices, San José, CA, USA). Acquired data were digitized at a sampling frequency of 20 kHz and filtered at 2 kHz with a Digidata 1322A (Axon Instruments, Palo Alto, CA, USA). Signals were acquired and analyzed offline with pCLAMP 10.6 software (Molecular Devices).

### Determination of intrinsic properties

The resting membrane potential (RMP) was determined after the initial membrane break-in from giga-seal to whole-cell configuration in voltage clamp mode. After 2–3 min. of stabilization, a series of negative and positive voltage commands from −100 to +50 (500 ms, 250 pA increments) were injected to determine the input resistance (R_N_) and membrane time constant (1_memb_). The R_N_ was calculated as the slope value of a first-order polynomial function f(*x*) = *mx* + *b* fitted to the current–voltage relationship near RMP. The 1_memb_ was determined by fitting a mono-exponential decay function to a voltage response elicited by injecting a negative current step (1 s, −30 pA). The membrane capacitance (C_m_) was calculated as the ratio of 1_memb_ to R_N_, as previously reported.

### Determination of macroscopic currents

Whole-cell voltage-clamp configuration was performed to evoke total membrane currents by the injection of depolarizing pulses (−90 to +60 mV, 10-mV steps, 500 ms) from a holding potential of −70 mV. The P/4 protocol was employed to subtract leak currents online, and the recording quality was monitored online by incorporating the following criteria: holding currents < 50 pA, stable RMP, and access resistance (< 15 MW, < 20% drift). To generate I-V plots, the current amplitude was measured either at the maximum peak amplitude or as the average over a 100-msec interval at the steady-state current.

### Statistical analysis

No statistical method was used to determine the sample size. The statistical analysis was performed with GraphPad Prism 8. The graphs represent the mean ± the standard error of the mean (SEM). The data normality was previously determined with a Kolmogorov-Smirnov test. Most of the data did not meet the normality criteria; and non-parametric tests were applied: Mann-Whitney U for comparing two experimental groups and Kruskal-Wallis for comparing more than two groups. A one-way analysis of variance (ANOVA) test was applied, followed by Tukey’s post-hoc test in cases where normal distribution was observed. In cases where the comparison was between two groups, the Student’s t-test was applied. In all cases, the accepted level of significance was set to *p* < 0.05 or a higher statistical significance.

## Results

### Progenitor markers and GABA_B_ receptor subunits are co-expressed in neural progenitor cells of the subventricular zone

Neurospheres derived from neural precursor cells of the subventricular zone (NPC SVZ) were obtained from CD1 mice on postnatal day eight and cultured in a DMEM/F12 medium supplemented with B27(1%), EGF (20 nM), and FGF (20 nM). The expression of progenitor and proliferation markers was then determined. Figures 1A and 1B show secondary neurospheres expressing the progenitor marker nestin and the proliferation marker Ki-67, thus confirming the progenitor origin of the cultured cells. Next, we analyzed if secondary neurospheres expressed the G-protein coupled receptor for gamma-aminobutyric acid receptor B (GABA_B_R). GABA_B_R is a metabotropic receptor assembled from two subunits: GABA_B1_ and GABA_B2_. Therefore, immunocytochemistry assays were individually performed against each receptor’s subunit. Figures 1C and 1D show the immunoreactivity of GABA_B1_ and GABA_B2_, respectively. The GABA_B_ subunits’ presence indicates that secondary neurospheres derived from NPC SVZ expressed the metabotropic GABA_B_ receptors in addition to the ionotropic GABA_A_ receptors (Stewart et al., 2002).

**Figure 1.**
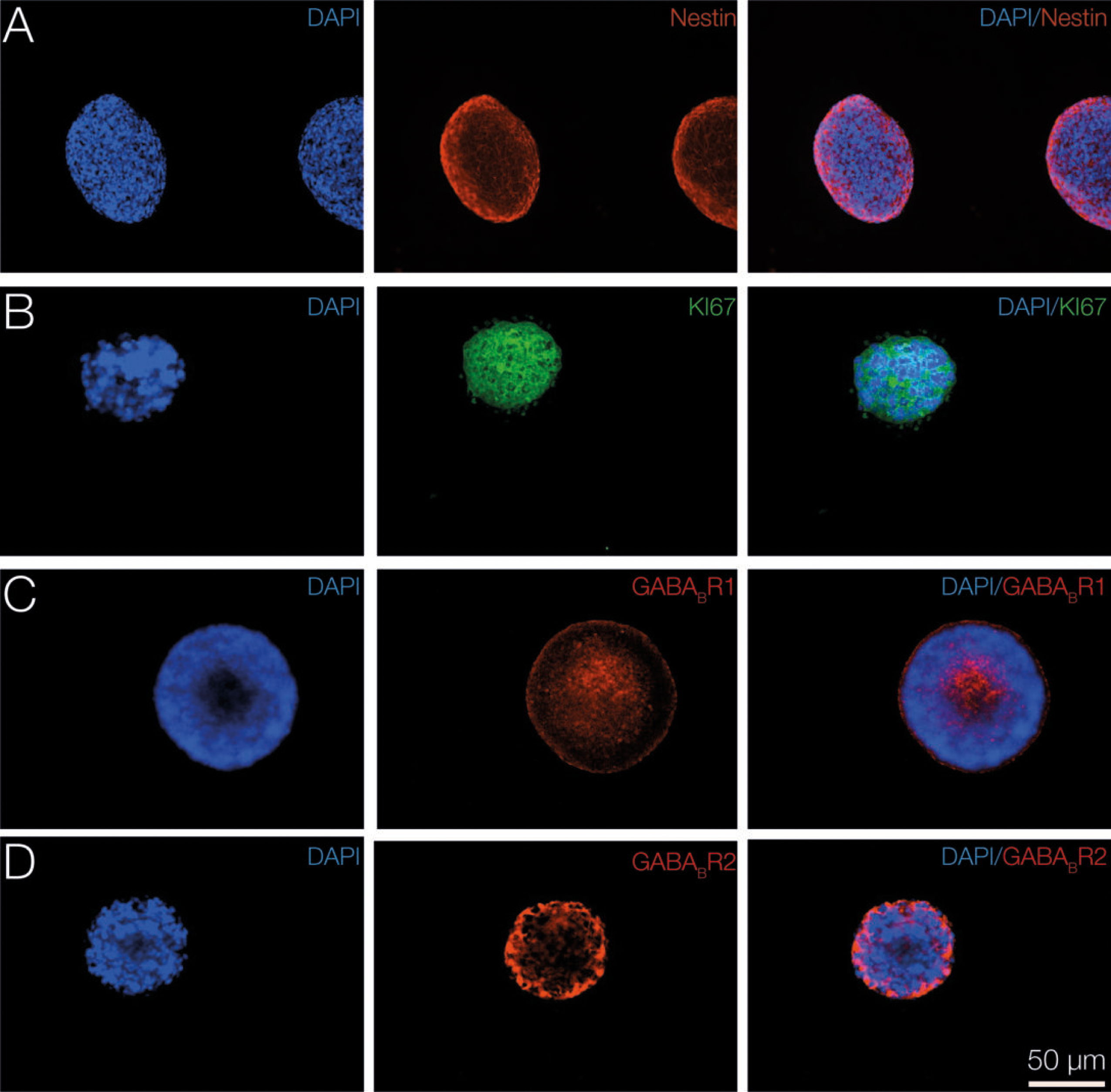
Expression of neural precursor cell markers and GABA_B_ receptor subunits in secondary neurospheres derived from the subventricular zone. Representative neurospheres derived from the subventricular zone. Secondary neurospheres exhibited a mean ratio of 70 ± 5 µM. The nuclei were stained with DAPI (blue). The neurospheres were immunopositive to **A)** nestin (red), **B)** KI67 (green), and the subunits of the GABA_B_ receptor **C)** GABA_B_R1 (red) and **D)** GABA_B_R2 (red). The microphotographs were taken with a 20x magnification lens. The scale bar applies to all panels.

### Activation of GABA_A_R with taurine induces neuronal differentiation and neurite outgrowth in NPC SVZ

We previously reported that disaggregated NPC SVZ incubated with taurine (14 days/10 mM) undergoes increased expression of neuronal markers accompanied by the development of passive and active electrophysiological properties (Gutierrez-Castañeda et al., 2023). The first observation was further investigated in the present study. The upper panels in Figure 2A show representative bright-field (10x) microphotographs of disaggregated NPC SVZ cultures in the control condition and incubated with taurine; the lower panels show immunofluorescence against doublecortin (DCX). Compared to the control condition, the taurine-treated cells exhibited an increased number of DCX+ cells (Mann Whitney test, U = 2; *p* < 0.01, Figure 2B) and increased development of somatic prolongations or neurites (Mann Whitney test, U = 426; *p* < 0.001; Figure 2C).

**Figure 2.**
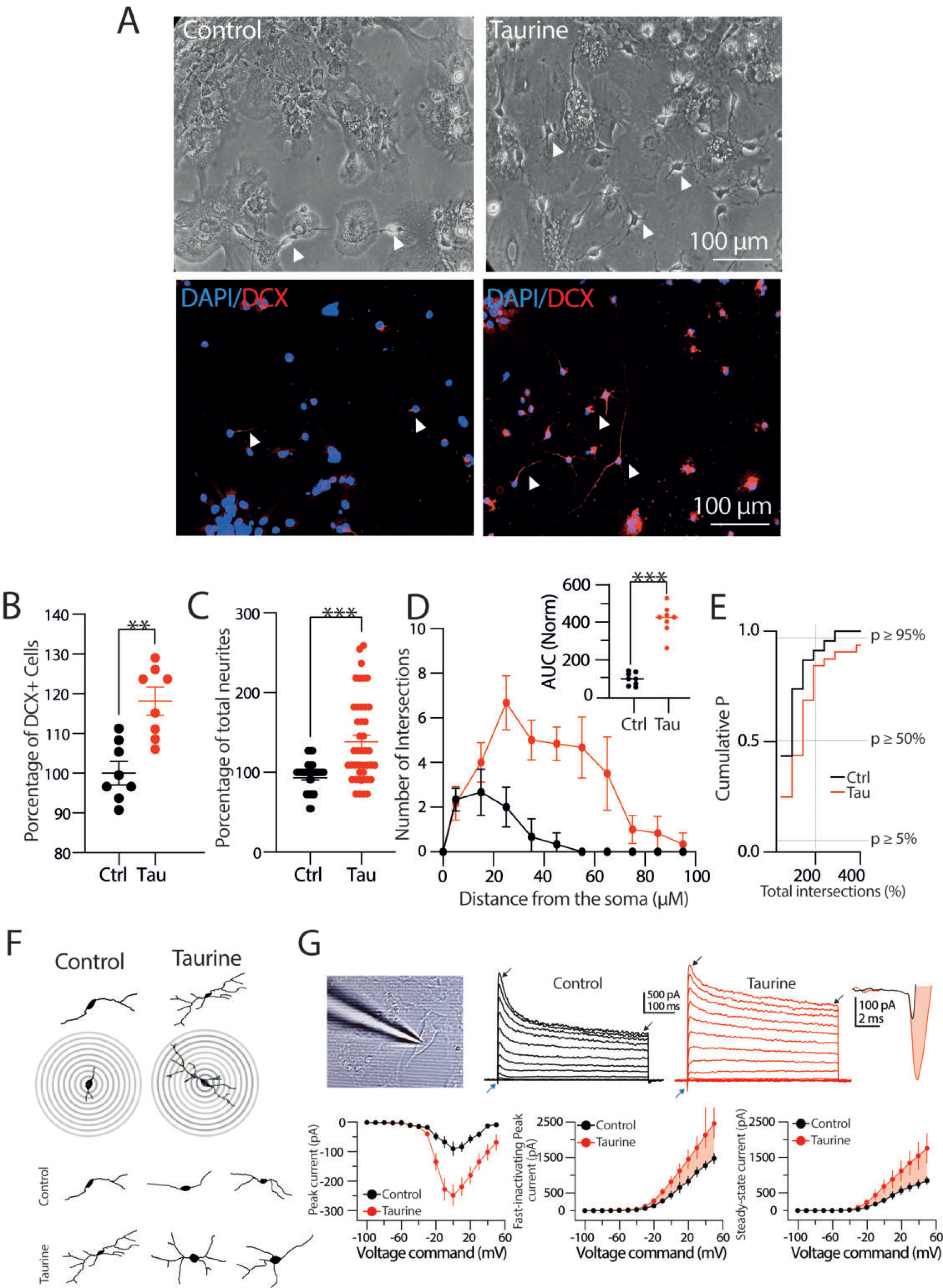
The interaction of the GABA_A_ receptor and taurine promotes neurite outgrowth of NPC SVZ. **A)** Representative bright-field and immunofluorescence assays microphotographs against DCX for immature neurons in the control condition or in taurine (10 mM). The nuclei were stained with DAPI (blue) and DCX (red). Arrowheads indicate cells with neuronal-type morphology used for the analyses included in this study. **B)** Scatter plot showing the percentage of DCX+ cells in the control and in taurine (*n* = 8 cell cultures for each experimental condition). **C)** Scatter plot with the total number of neurites in the control condition (*n* = 36) and in taurine (*n* = 40 DCX+ cells). **D)** Scatter plot of the Sholl-like analysis contrasting the number of neurite intersections in the control condition and the taurine-treated cells. Upper inset: area under the curve (AUC) chart showing the total area obtained from the Sholl-like analysis performed in the control and taurine-treated cells (*n* = 8 DCX+ cells). **E)** Cumulative probability chart summarizing the number of neurite intersections in the control cells versus the taurine-treated cells. The right shift of the taurine-treated cells mirrors the increased number of neurite intersections (*n* = 8 for each experimental condition). **F)** Upper panels: representative digital reconstructions and concentric rings (distance between rings = 10 µM) used for the Sholl-like morphometric analysis. Bottom panel: examples of the increased number of primary, secondary, and tertiary neurites of the taurine-treated cells. **G)** Left panel: a patch-clamp pipette attached to an NPC SVZ incubated with taurine. Notice the cell body and the neurite outgrowths. Middle panel: representative current traces obtained in a control and a taurine-treated cell. The blue arrowheads indicate the inward current, the black arrowheads indicate the fast-inactivating and the persistent, steady-state outward current. Right panel: magnification of the inward currents, black trace control, red trace, inward current of a taurine-treated cell. Bottom left panel: scatter plot of the voltage dependence and maximal amplitude of the inward current. Middle panel: scatter plot showing the maximal peak amplitude of the fast-inactivating outward current. Right panel: scatter plot depicting the amplitude of the non-inactivating steady-state outward current. \*\**p* < 0.01; \*\*\**p* < 0.001 or higher statistical significance. Error bars indicate SEM.

Next, a Sholl-like morphometric analysis was used to determine the pattern of neurite branching of the disaggregated cells treated with taurine. Digital reconstructions were performed from the bright-field microphotographs, and the resulting images were processed to trace neuronal arborizations. Figure 2F shows representative digital reconstructions of NPC SVZ obtained in both experimental conditions and the overlapped concentric circles (distance = 10 µM) from which the values of the Sholl-like plot were obtained. The taurine-treated cells exhibited an increased number of neurite intersections compared to the control cells (Figures 2D and 2F). The increased number of intersections is summarized in the inset graph in Figure 2D, showing the area under the curve (AUC) obtained from the Sholl scatter plot (AUC in taurine-treated cells = 310.4 ± 39.4% compared with control AUC; t-test; *p* < 0.001). The cumulative probability chart in Figure 2E shows the distribution of the total number of intersections in the control cells and cells treated with taurine. Morphologically, the control cells displayed bipolar-oriented somata with one or two primary neurites; by contrast, the taurine-treated cells exhibited a multipolar morphology with extensive neuritic arborizations and complex branching. Likewise, the number of neurites and intersections was significantly higher in the proximity of the somata of the taurine-treated cells (see representative examples in Figure 2F).

Next, we examined the functional expression of macroscopic currents expressed by the NPC SVZ using the whole-cell patch-clamp technique. Under the voltage-clamp mode, we applied a protocol (see methods for details) that sequentially elicited an inward current followed by an outward current. Taurine incubation dramatically increased the inward current compared to the control cells (inward current in the control condition = −76.2 ± 24 pA; in the taurine-treated cells = −248.1 ± 36.3 pA; t-test; p < 0.01; Figure 2G; blue arrowheads and far-right traces). On the other hand, the outward current exhibited two kinetic components: a fast-inactivating current peak followed by a steady, non-inactivating current that resembles the neuronal *I*_A_ and the *I*_D_ potassium currents (See Griego et al., 2021, 2022). Interestingly, the taurine-treated cells exhibited an increased amplitude of the fast-inactivating outward current (fast-inactivating current in the control condition = 1,474.3 ± 150 pA; in the taurine-treated cells = 2,621.4 ± 42 pA; t-test; p < 0.05; Figure 2G; black arrowheads in the middle panel current traces) and the steady, non-inactivating current (steady-state outward current in the control condition = 847 ± 111 pA; in the taurine-treated cells = 1,957.5 ± 421 pA; t-test; p < 0.05; Figure 2G; black arrowheads in the middle panel current traces). Interestingly, the somatic input resistance (R_N_) for the taurine-treated cells dropped ≈43% compared to the control cells (R_N_ in the control condition = 452.3 ± 55.49 MΩ; in the taurine-treated cells = 196.2 ± 17 MΩ; t-test; *p* < 0.001). Lastly, the relationship membrane time-constant / somatic input resistance ratio (1_memb_/R_N_) yielded a membrane capacitance (C_m_) of 47.39 ± 9.22 pF for the control and 41.47 ± 14.56 pF (*ns*) for the taurine-treated cells, suggesting that the observed increase in the current amplitude was unrelated to changes in the somatic area.

Next, we corroborated that the pharmacological blockade of GABA_A_R decreases the neuronal differentiation process mediated by taurine (see also Gutierrez-Castañeda et al., 2023). Figure 3A shows bright-field microphotographs and the respective immunofluorescence assay against DCX performed in the control, the taurine-, and the taurine + picrotoxin (PTX, 100 µM)–treated cells. Figure 3B shows the decreased number of DCX+ cells in the presence of PTX (DCX+ cells in the presence of taurine = 118.1 ± 3.5%; one-way ANOVA; Tukey’s test; *p* < 0.01; in the presence of taurine + PTX = 72.4 ± 3.4%; one-way ANOVA; Tukey’s test; *p* < 0.001). Consistent with this finding, Figure 3C shows a reduced number of neurites in the taurine + PTX condition (total number of neurites in the taurine-treated cells compared to the control cells = 138.3 ± 8.1%; one-way ANOVA; Tukey’s test; *p* < 0.001; in taurine + PTX vs. the taurine-treated cells = 52.6 ± 7.6%; one-way ANOVA; Tukey’s test; *p* < 0.001). Likewise, the blockade of GABA_A_R with PTX substantially decreased the intersections computed in the Sholl-like analysis (Figures 3D and 3F). This finding is summarized in the inset AUC graph (AUC in the presence of taurine = 310.4 ± 39.42%; one-way ANOVA; Tukey’s test; *p* < 0.001; AUC in the presence of taurine + PTX = 81.25 ± 33.74 %; one-way ANOVA; Tukey’s test; *p* < 0.001). The cumulative probability chart in Figure 3E shows the distribution of the total number of intersections in the three experimental conditions. Representative examples of the decreased neurite branching in response to the blockade of GABA_A_R with PTX are shown in the digital reconstructions (Figure 3F). The whole-cell patch clamp recordings revealed a series of changes in the amplitude of the macroscopic currents when GABA_A_R was blocked. Incubation of taurine + PTX decreased the maximal amplitude of the inward current compared to the taurine-treated cells (inward current in the taurine-treated cells = −248.09 ± 36.30 pA; in the taurine + PTX–treated cells = −32.42 ± 11.66 pA t-test; *p* < 0.001; Figure 3G; blue arrowheads and far-right traces). On the other hand, both the fast-inactivating and the steady-state outward currents exhibited a decreased amplitude in the presence of PTX (fast-inactivating current in the taurine-treated cells = 2,154.5 ± 422.03 pA; in the presence of taurine + PTX = 1,057.1 ± 129.8 pA. t-test; *p* < 0.01; steady-state outward current in the taurine-treated cells = 1,557.5 ± 421 pA; in the presence of taurine + PTX = 617.45 ± 106.6 pA. t-test; *p* < 0.01; Figure 3G; white arrowheads in the middle panel current traces). In the taurine + PTX–treated cells, the R_N_ did not exhibit a significant change compared to the control cells (R_N_ in the taurine-treated cells = 204.6 ± 18.2 MΩ; in taurine + PTX = 513.5 ± 114 MΩ; t-test; *p* < 0.01). No changes were found in the C_m_ of the taurine + PTX–treated cells compared to the taurine-treated cells (C_m_ in the taurine-treated cells = 47.39 ± 9.22 pF; in taurine + PTX = 55.10 ± 18.13 pF). Collectively, these experiments indicate that activation of the GABA_A_R is necessary for the morphogenic process, neurite development, and functional expression of ion channels mediated by taurine.

**Figure 3.**
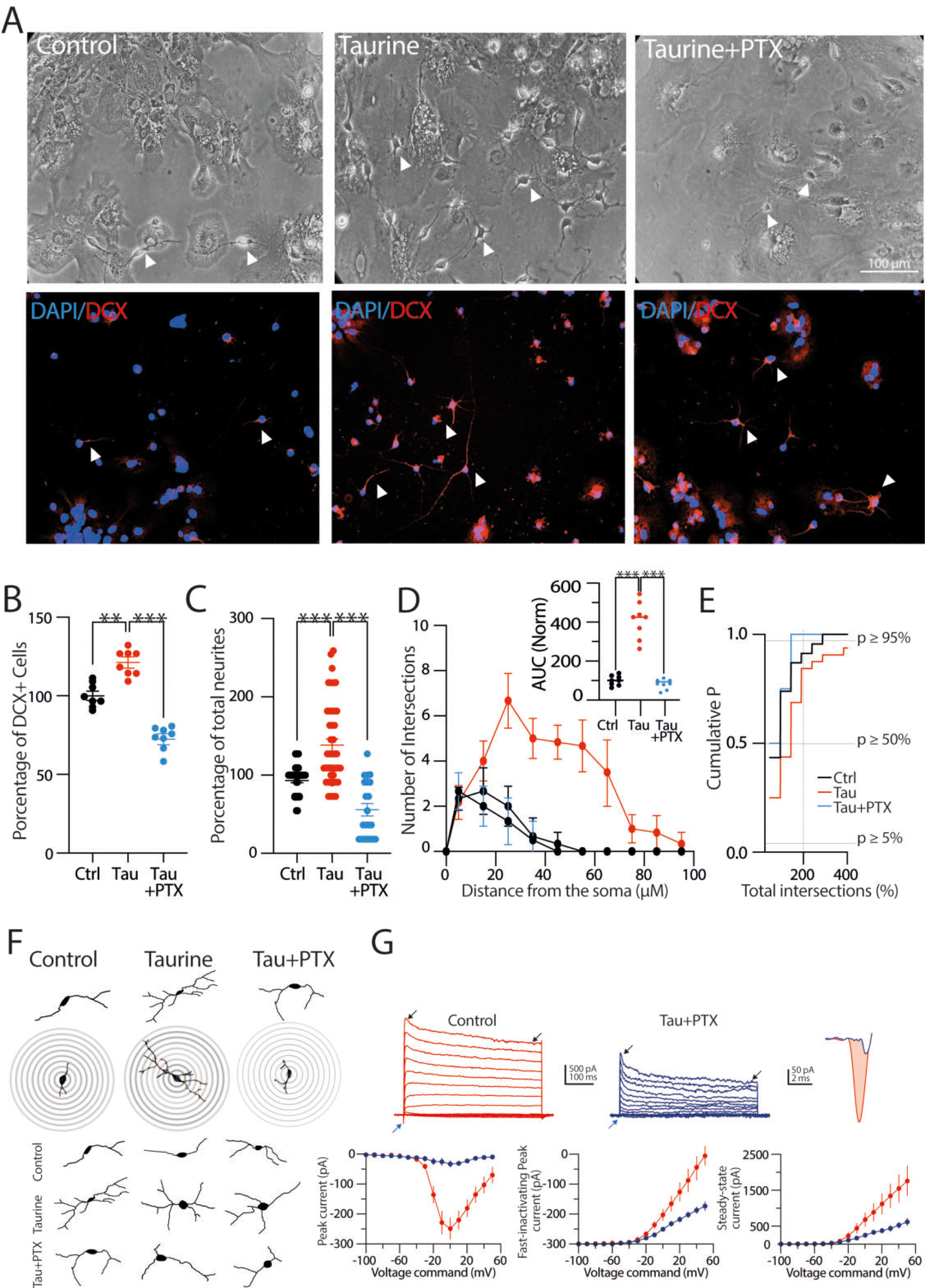
Pharmacological blockade of GABA_A_ receptor inhibits neurite outgrowth of NPC SVZ. **A)** Bright-field and immunofluorescence assays. Nuclei were stained with DAPI (blue) and DCX (red). Arrowheads show cells with a neuronal-type morphology obtained in the indicated experimental conditions. **B)** Scatter plot summarizing the percentage of DCX+ cells in the control and in the presence of taurine and taurine + PTX (*n* = 8 cell cultures for each experimental condition). **C)** Scatter plot showing the total number of neurites in the indicated experimental conditions. PTX prevented the neurite outgrowth observed stimulated with taurine (control *n* = 34 cells; taurine *n* = 38 cells; taurine + PTX *n* = 27 DCX+ cells). **D)** Sholl-like analysis with the number of neurite intersections in the indicated experimental conditions. Upper inset: AUC chart summarizing the findings of the Sholl-like analysis (*n* = 8 DCX+ cells). **E)** Cumulative probability chart. The left shift of the taurine + PTX–treated cells (blue line) mirrors the decreased number of neurite intersections when GABA_A_ receptors were blocked (control *n* = 34 cells; taurine *n* = 38 cells; taurine + PTX *n* = 27 DCX+ cells). **F)** Representative digital reconstructions and concentric rings used for the Sholl-like analysis. Notice the decreased number of primary, secondary, and tertiary neurites of the taurine + PTX–treated cells (right panel). **G)** Upper panel: representative current traces obtained from cells treated with taurine or taurine + PTX. Blue arrowheads indicate the inward currents, black arrowheads the fast-inactivating and the persistent, steady-state outward current. Upper right panel: magnification of the inward current from a taurine-treated cell (red trace) and a taurine + PTX–treated cell (blue trace). Bottom left panel: voltage dependence and maximal amplitude of the inward current. Middle panel: the maximal amplitude of the fast-inactivating outward current. Right panel: the steady-state amplitude of the non-inactivating outward current. ns = non-significant statistical difference; \*\**p* < 0.01; \*\*\**p* < 0.001 or higher statistical significance. Error bars indicate SEM.

### Activation of GABA_B_ receptors does not stimulate the differentiation process of NPC SVZ

Because secondary neurospheres express both subunits of the GABA_B_R (see Figures 1C and 1D), we next determined the role of the metabotropic GABA_B_R in the differentiation process of NPC SVZ. The cultures were exposed to the GABA_B_R agonist baclofen (100 μM) in the presence of taurine (10 mM/14 days), and the expression of DCX was evaluated. Figure 4A shows bright-field microphotographs and the respective immunofluorescence assays against DCX in the control, the taurine-, and the taurine + baclofen–treated cells. Activation of GABA_B_R with baclofen + taurine did not modify the number of DCX+ cells compared to the taurine-treated cells (number of DCX+ cells in the presence of taurine compared to the control cells = 118.1 ± 3.5%; one-way ANOVA; Tukey’s test; *p* < 0.01; in the presence of taurine + baclofen compared with the taurine-treated cells = 110 ± 4.4%; ns; Figure 4B). Moreover, the activation of GABA_B_R with baclofen decreased the number of neurites (total number of neurites in taurine compared to the control cells = 138.3 ± 8.1%; one-way ANOVA; Tukey’s test; *p* < 0.01; in taurine + baclofen compared with the taurine-treated cells = 83.1 ± 7.7%; one-way ANOVA; Tukey’s test; *p* < 0.001; Figure 4C). Likewise, the neurite intersections were also decreased in the taurine + baclofen condition, as shown in Figure 4D and the inset and the concentric circles of the Sholl-like plot analysis in Figure 4F (AUC in the presence of taurine compared to the control cells = 310.1 ± 40%; one-way ANOVA; Tukey’s test; *p* < 0.001; AUC in the presence of taurine + baclofen compared with the taurine-treated cells = 189.6 ± 37.5 %; one-way ANOVA; Tukey’s test; *p* < 0.001).

**Figure 4.**
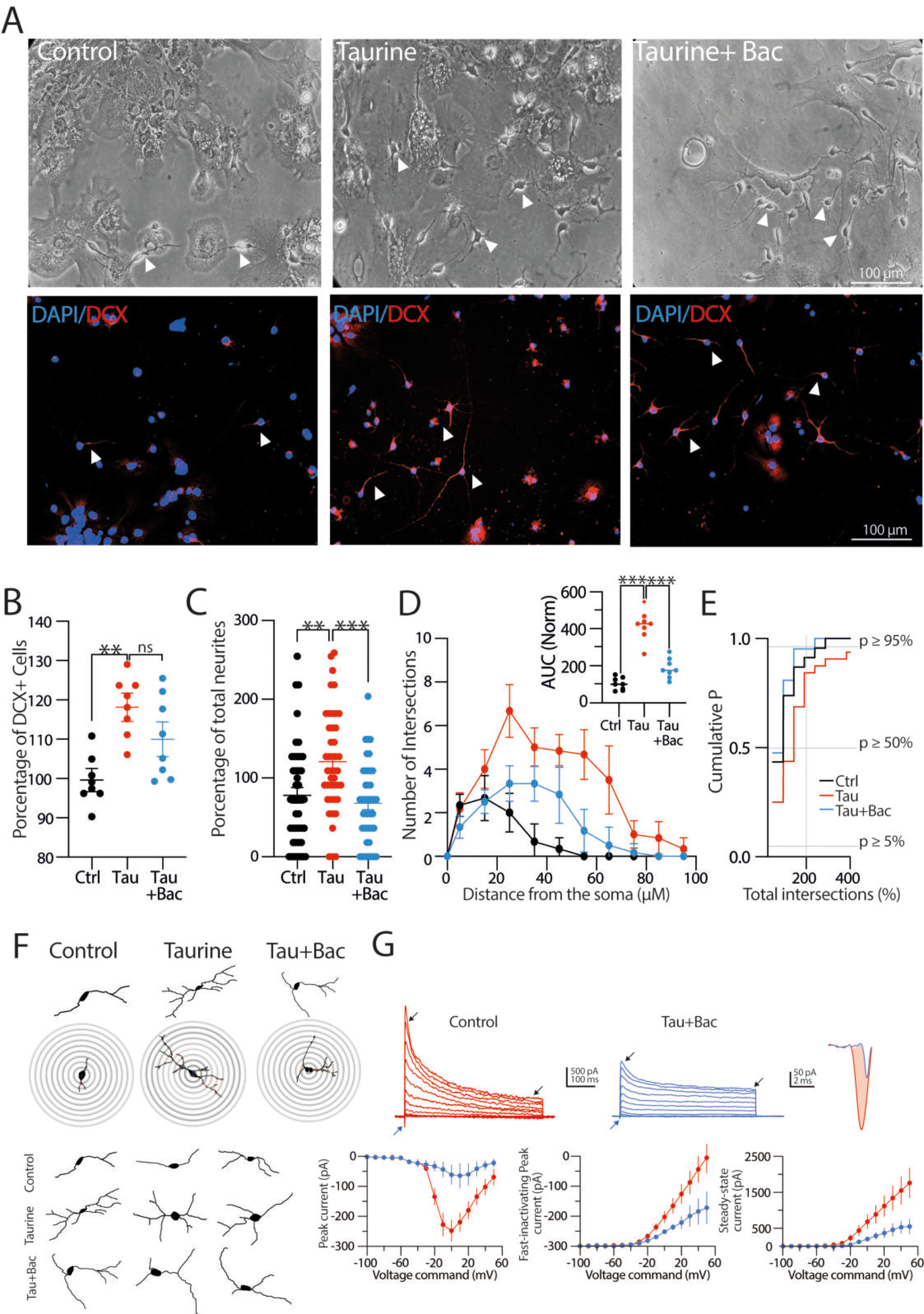
Activation of the GABA_B_ receptor with baclofen does not stimulate the differentiation process of NPC SVZ. **A)** Bright-field and immunofluorescence assays microphotographs for DCX in the control cells, taurine-, and taurine + baclofen (100 µM)–treated cells. Nuclei were stained with DAPI (blue) and DCX (red). Arrowheads show cells with a neuronal-type morphology obtained in the indicated conditions. **B)** Scatter plot showing the percentage of DCX+ cells in the indicated experimental conditions. Stimulation with baclofen did not alter the number of DCX+ cells (*n* = 8 cell cultures for each experimental condition). **C)** Comparison of the total number of neurites in the indicated experimental conditions. The combination of taurine + baclofen did not stimulate neurite outgrowths compared to the taurine-treated cells (control *n* = 33 cells; taurine *n* = 43 cells; taurine + baclofen *n* = 37 DCX+ cells). **D)** Sholl-like analysis obtained from the indicated experimental conditions. Upper inset: AUC chart summarizing the findings of the Sholl-like analysis (*n* = 8 DCX+ cells). **E)** Cumulative probability chart of the number of neurite intersections in the three experimental conditions (control *n* = 33 cells; taurine *n* = 43 cells; taurine + baclofen *n* = 37 DCX+ cells). **F)** Digital reconstructions and concentric rings used for morphometric analyses. Notice the subtle changes in the number of primary, secondary, and tertiary neurites of the taurine + baclofen–treated cells (right panel). **G)** Upper panel: representative current traces obtained from cells treated with taurine or taurine + baclofen. Blue arrowheads indicate the inward currents, black arrowheads the fast-inactivating and the persistent, steady-state outward current. Upper right panel: magnification of the inward current from a taurine-treated cell (red trace) and taurine + baclofen–treated cell (blue trace). Bottom left panel: voltage dependence and maximal amplitude of the inward current. Middle panel: the maximal amplitude of the fast-inactivating outward current. Right panel: the steady-state amplitude of the non-inactivating outward current. ns = non-statistical significance; \*\**p* < 0.01; \*\*\**p* < 0.001 or higher statistical significance. Error bars indicate SEM.

The cumulative probability chart in Figure 4E shows the distribution of the total number of intersections in the three experimental conditions. Morphologically, the cells treated with taurine + baclofen exhibited similar characteristics to the control cells, including short and few neurite outgrowths (Figure 4F). The lack of action of the GABA_B_R stimulation was mirrored in the amplitude of the inward current (inward current in the taurine-treated cells = −248.09 ± 36.30 pA; in the taurine + baclofen– treated cells = −64.99 ± 41.94 pA; t-test; *p* < 0.01; Figure 4G; blue arrowheads and far-right traces) and outward current’s components (fast-inactivating current in the taurine-treated cells = 2,154.5 ± 422 pA; in the presence of taurine + baclofen = 1,070 ± 450 pA. t-test; (*ns*); steady-state outward current in the taurine-treated cells = 1,557.5 ± 421 pA; in the presence of taurine + baclofen = 550 ± 207 pA. t-test; (*ns*); Figure 4G; white arrowheads in the middle panel current traces). In the presence of tau + baclofen, the R_N_ showed a significant change compared to the taurine-treated cells (R_N_ in the taurine-treated cells = 190.8 ± 19.7 MΩ; in the taurine + baclofen–treated cells = 463.3 ± 66 MΩ; t-test; *p* < 0.001).

No changes were found in the C_m_ of the taurine + baclofen–treated cells compared to taurine-treated cells (C_m_ in the taurine-treated cells = 48.1 ± 9.22 pF; in taurine + baclofen = 43.84 ± 11.42 pF). Collectively, these results demonstrate that baclofen-induced GABA_B_R activation does not contribute to the neuronal differentiation process of the NPC SVZ or to the balanced expression of macroscopic currents; on the contrary, GABA_B_R activation appears to interfere with neurite outgrowth and the functional expression of ion channels.

### Blockade of the GABA_B_ receptors favors the neuronal differentiation process, complexity and the number of neurites of NPC SVZ

Next, we explored the effects of the blockade of GABA_B_R in the differentiation process. NPC SVZ were exposed to the selective GABA_B_R antagonist CGP 55845 (5 μM) in the presence of taurine (10 mM/14 days), and the expression of DCX was evaluated. Bright-field microphotographs and immunofluorescence assays against DCX performed in the control, taurine-, and taurine + CGP 55845–treated cells are shown in Figure 5A. Strikingly, the blockade of GABA_B_R with CGP 55845 increased the number of DCX+ cells by a similar amount to the observed with taurine (DCX+ cells in the presence of taurine compared to the control cells = 118.1 ± 3.5%; one-way ANOVA; Tukey’s test; *p* < 0.01; in the presence of CGP 55845 + taurine compared to the taurine-treated cells = 116.8 ± 2.5%; one-way ANOVA; Tukey’s test; *p* < 0.05. Figure 5B). Likewise, these cells exhibited a higher number of neurites (total number of neurites in the presence of taurine compared to the control cells = 138.3 ± 8%; one-way ANOVA; Tukey’s test; *p* < 0.001; in the presence of taurine + CGP 55845 compared to the taurine-treated cells = 127.2 ± 8.6; one-way ANOVA; Tukey’s test; ns; Figure 5C). Additionally, the blockade of GABA_B_R increased the number of neurite intersections (Figure 5D and the concentric circles of the Sholl-like plot in Figure 5F). The increased intersection number in response to CGP 55845 + taurine is summarized in the inset graph (AUC in the presence of taurine compared to the control cells = 310.4 ± 40%; one-way ANOVA; Tukey’s test; *p* < 0.001; AUC in the presence of taurine + CGP 55845 compared to the taurine-treated cells = 516.7 ± 45.55%; one-way ANOVA; Tukey’s test; *p* < 0.001). The cumulative probability chart in Figure 5E shows the distribution of the total number of intersections in the control, in cells treated with taurine, and in cells treated with taurine + CGP 55845. Morphologically, the neurites of the CGP 55845–treated cells exhibited a robust neuronal-like morphology, with radiated neurite outgrowth and a more complex ramification pattern than those observed in the taurine-treated group. Interestingly, the amplitude of the inward currents did not increase in the presence of CGP 55845 (inward current in the taurine-treated cells = −248.09 ± 36.30 pA; in the taurine + CGP 55845–treated cells = −29.78 ± 2.84 pA; t-test; *p* < 0.01; Figure 5G; blue arrowheads and far-right traces). On the other hand, the amplitude of the fast-inactivating, and the persistent current increased in the presence of CGP 55845 (fast-inactivating current in the taurine-treated cells = 2,050.4 ± 142 pA; in the presence of taurine + CGP 55845 = 978.41 ± 181 pA. t-test; *p* < 0.01; steady-state outward current in the taurine-treated cells = 1,559 ± 421 pA; in the presence of taurine + baclofen = 1,219.26 ± 171 pA; t-test, *ns*; Figure 4G). Regarding the R_N_, the tau + CGP 55845 treatment caused a slight drop (≈19%) compared to the control cells (R_N_ in the control condition = 465 ± 55 ΩOhm; in the taurine + CGP 55845– treated cells = 379.5 ± 52 ΩOhm; t-test; *p* < 0.05). No changes were found in the Cm of the cells treated with CGP 55845 (C_m_ in the taurine-treated cells = 47.39 ± 9.22 pF; in taurine + baclofen = 53.48 ± 12.1 pF). These findings demonstrate that the blockade of the metabotropic GABA_B_R enhances the morphogenic process of NPC SVZ with limited effects on the functional expression of macroscopic currents. The latter finding requires additional investigation.

**Figure 5.**
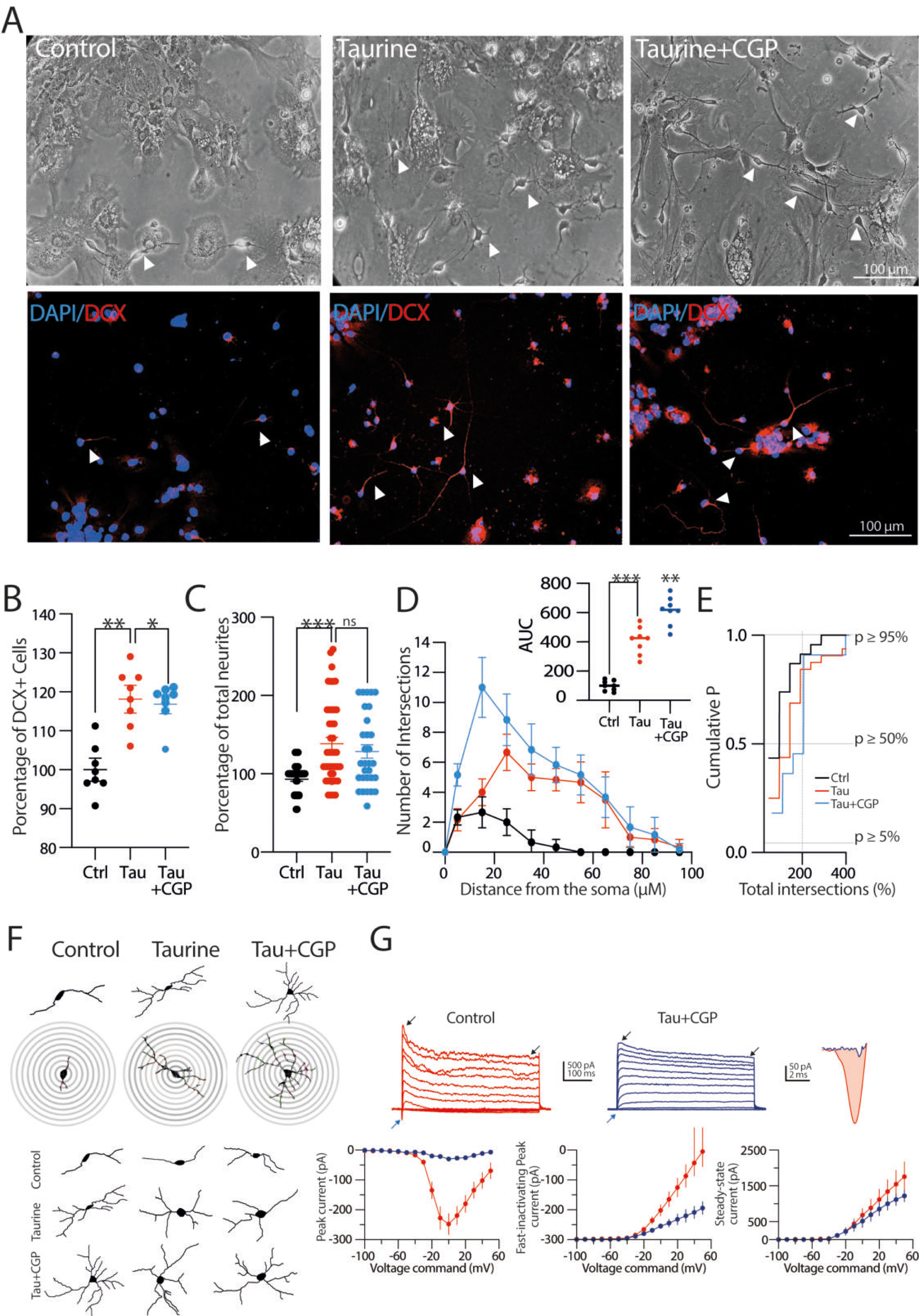
The blockade of GABA_B_ receptors with CGP 55845 favors the differentiation process and neurite outgrowth of NPC SVZ. **A)** Bright-field and immunofluorescence assays microphotographs for DCX in the control condition, taurine, and taurine + CGP 55845 (5 µM). Arrowheads indicate cells with a neuronal-type morphology. **B)** Scatter plot summarizing the percentage of DCX+ cells in the indicated experimental conditions (*n* = 8 cell cultures for each experimental condition). **C)** Scatter plot summarizing the total number of neurites. Taurine + CGP 55845 increased the total number of neurites (control *n* = 38 cells; taurine *n* = 47 cells; taurine + CGP 55845 *n* = 43 DCX+ cells). **D)** Sholl-like plot contrasting the number of intersections in the indicated experimental conditions. Upper inset: AUC chart (*n* = 8 cells for each experimental condition). **E)** Cumulative probability chart summarizing the number of neurite intersections. The right shift of the taurine + CGP 55845–treated cells mirrors the increased number of neurite intersections (*n* = 6 DCX+ cells). **F)** Representative digital reconstructions and concentric rings. Notice the increased number of primary, secondary, and tertiary neurites of the taurine + CGP 55845–treated cells. **G)** Upper panel: representative current traces obtained from NPC SVZ treated with taurine or taurine + CGP 55845. Blue arrowheads indicate the inward currents, black arrowheads the fast-inactivating and the persistent, steady-state outward current. Upper right panel: magnification of the inward current from a taurine-treated cell (red trace) and a taurine + CGP 55845– treated cell (blue trace). Bottom left panel: voltage dependence and maximal amplitude of the inward current. Middle panel: the maximal amplitude of the fast-inactivating outward current. Right panel: the steady-state amplitude of the non-inactivating outward current. ns = non-statistical significance; \**p* < 0.05; \*\**p* < 0.01; \*\*\**p* < 0.001 or higher statistical significance. Error bars indicate SEM.

### A rise in intracellular calcium is necessary for the neuronal differentiation process mediated by taurine

Because taurine is a well-known modulator of intracellular Ca^2+^ homeostasis (Wu and Prentice, 2010) we explored if calcium mobilization is involved in the morphogenic process of NPC SVZ mediated by taurine. To test this hypothesis, the cultures were preincubated with taurine + BAPTA-AM. As expected, the intracellular Ca^2+^ chelation with BAPTA-AM significantly reduced the number of neurites mediated by taurine (total number of neurites in the presence of taurine compared to the control cells = 138.3 ± 7.5%; *p* < 0.001; in the presence of taurine + BAPTA-AM compared to the taurine-treated cells = 35.16 ± 7.6%. *p* < 0.001; one-way ANOVA., Tukey’s test; Figure 6A). Likewise, BAPTA-AM decreased the total length of the neurites of the taurine-treated cells (total length of neurites in the presence of taurine = 27.4 ± 1.5 µm; *p* < 0.001; in the presence of taurine + BAPTA-AM = 11.5 ± 2 µm; *p* < 0.001. one-way ANOVA; Tukey’s test; Figure 6B). The effects of BAPTA-AM on the neurite outgrowth are shown in the Sholl-like plots (Figure 6C) and summarized in the AUC graph (AUC in the presence of taurine compared to the control cells = 410.4 ± 24.6%; *p* < 0.001; in the presence of taurine + BAPTA-AM = 107.1 ± 24%; Figure 6D).

**Figure 6.**
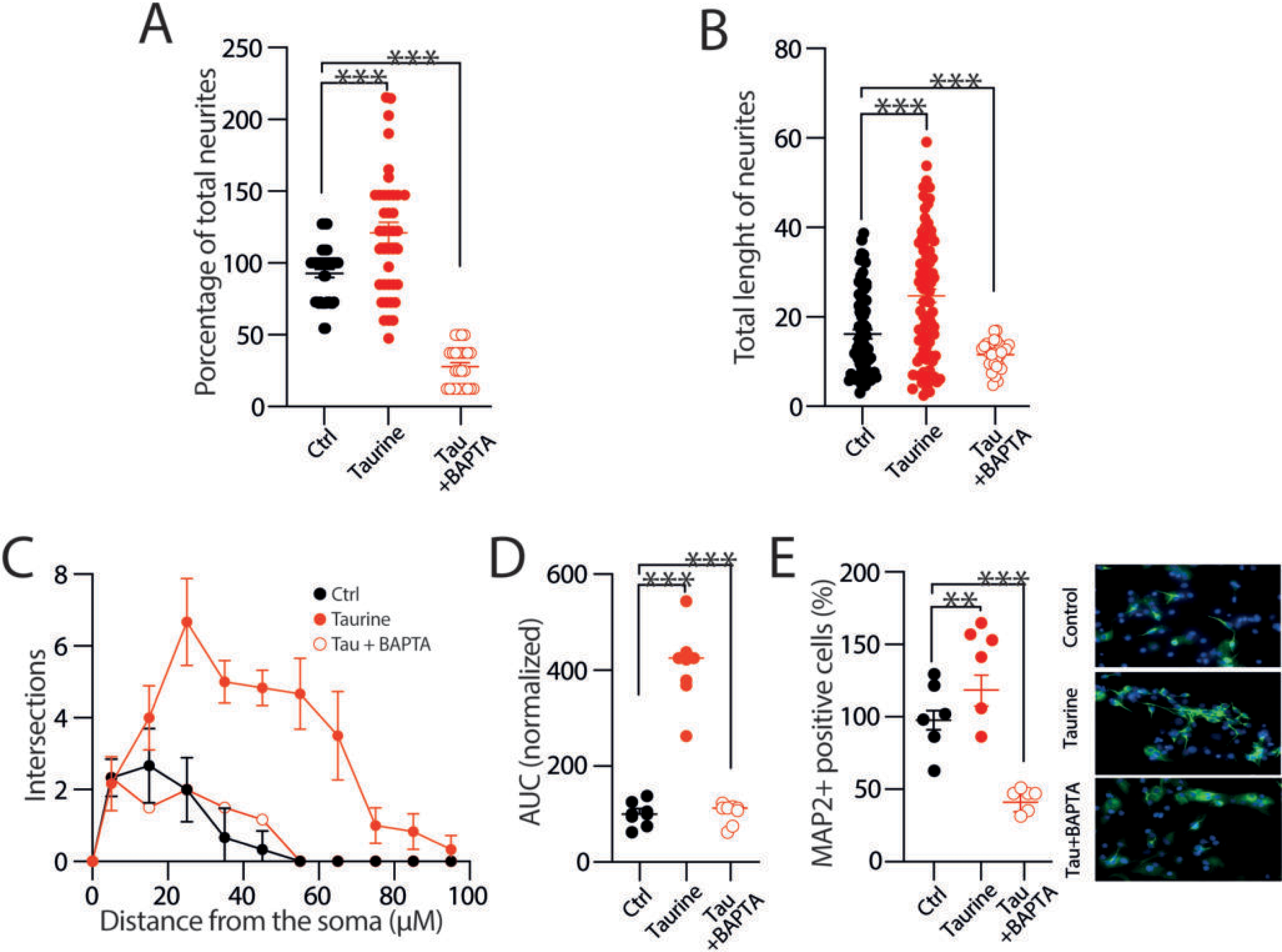
Taurine involves intracellular calcium mobilization for the differentiation process and neurite outgrowth of NPC SVZ. **A)** Scatter plot showing the number of neurites of NPC SVZ exposed to taurine or taurine + BAPTA-AM. Intracellular calcium chelation prevented neurite outgrowth. **B)** Scatter plot showing neurite length in the indicated experimental conditions. BAPTA incubation inhibited neurite outgrowth stimulated with taurine (control *n* = 31 cells; taurine *n* = 36 cells; taurine + BAPTA-AM *n* = 12 MAP2+ cells). **C)** Sholl-like plot contrasting the number of intersections in the indicated experimental conditions. **D)** AUC chart summarizing the findings of the Sholl-like analysis (*n* = 8 MAP2+ cells). **E)** Immunofluorescence assay microphotographs against MAP2 in taurine- and taurine + BAPTA–treated cells. BAPTA treatment reduced immunoreactivity to MAP2 (*n* = 6 cells for each experimental condition). ns = non-statistical significance; \*\**p* < 0.01; \*\*\**p* < 0.001 or higher statistical significance. Error bars indicate SEM.

We also performed an immunoreactivity assay against the microtubule-associated protein 2 (MAP2) in the cultures exposed to taurine + BAPTA-AM. Immunoreactivity to MAP2 is a strong indicator of stable neuronal morphology during neuronal maturation. In addition to affecting neurite outgrowth, we hypothesized that Ca^2+^ chelation with BAPTA would negatively affect neuronal maturation. As expected, BAPTA-AM decreased the MAP2 immunoreactivity (MAP2+ immunoreactivity in the presence of taurine compared to the control cells = 134.6 ± 13.3; *p* < 0.01; in the presence of taurine + BAPTA-AM = 69.90 ± 8.31%; *p* < 0.001; one-way ANOVA; Tukey’s test; Figure 6E). Collectively, these experiments demonstrate that the differentiation and maturation process of NPC SVZ mediated by taurine involves intracellular calcium mobilization.

To culminate this study, we investigated the role of Ca^2+^-dependent signaling cascades. The cultures were preincubated with taurine combined with the calcium– calmodulin (CaM)–dependent protein kinase II (CaMKII) inhibitor KN93, the extracellular-signal-regulated kinase (ERK1/2) inhibitor FR1805, and the Src-family protein-tyrosine kinase inhibitor Srcl. The blockade of CaMKII, ERK1/2, and Srcl inhibited the formation of neurites stimulated with taurine (total number of neurites in the presence of taurine compared to the control cells = 138.14 ± 6.4 μM; *p* < 0.001; in the presence of taurine + KN93 = 36.1 ± 8.1%; *p* < 0.01; in the presence of taurine + FR1805 = 38 ± 8.7%; *p* < 0.01; in the presence of taurine + Srcl = 44.6 ± 9%; *p* < 0.01; one-way ANOVA; Tukey’s test; Figure 7A) and inhibited the increase in their length (total length of neurites in the presence of taurine compared to the control cells = 24.7 ± 1.6 μM; *p* < 0.001; in the presence of taurine + KN93 = 14.4 ± 4 μM; *p* < 0.05; in the presence of taurine + FR1805 = 16.7 ± 2.2 μM; *p* < 0.05; in the presence of taurine + SrcI = 12.8 ± 2.5 μM; *p* < 0.01; one-way ANOVA; Tukey’s test; Figure 7B). Likewise, the inhibition of the signaling cascades impacted the number of intersections plotted in the Sholl-like analysis (Figure 7C) and the resulting AUC (AUC in the presence of taurine compared to the control cells = 410.4 ± 24.6%; *p* < 0.001; in the presence of taurine + FR1805 = 95.9 ± 24.6%; *p* < 0.001; in the presence of taurine + SrcI = 106.3 ± 24.6%; *p* < 0.001; one-way ANOVA; Tukey’s test; Figure 7D). Consistent with these findings, the immunoreactivity of MAP2 kinase was dramatically decreased when these signaling cascades were blocked (MAP2+ cells in the presence of taurine compared to the control cells = 134.6 ± 13.3%; *p* < 0.001; in the presence of taurine + KN93 = 61.6 ± 12%; *p* < 0.001; in the presence of taurine + FR1805 = 57.5 ± 13.3%; *p* < 0.001; in the presence of taurine + SrcI = 52.3 ± 13.3%; *p* < 0.001; one-way ANOVA; Tukey’s test; Figure 7E). Collectively, these results demonstrate that CaMKII, ERK1/2, and the Src kinase play a crucial role in the taurine-mediated neuronal differentiation process. More specifically, inhibition of these signaling cascades negatively impacts the morphology and complexity of the differentiation process of the NPC SVZ.

**Figure 7.**
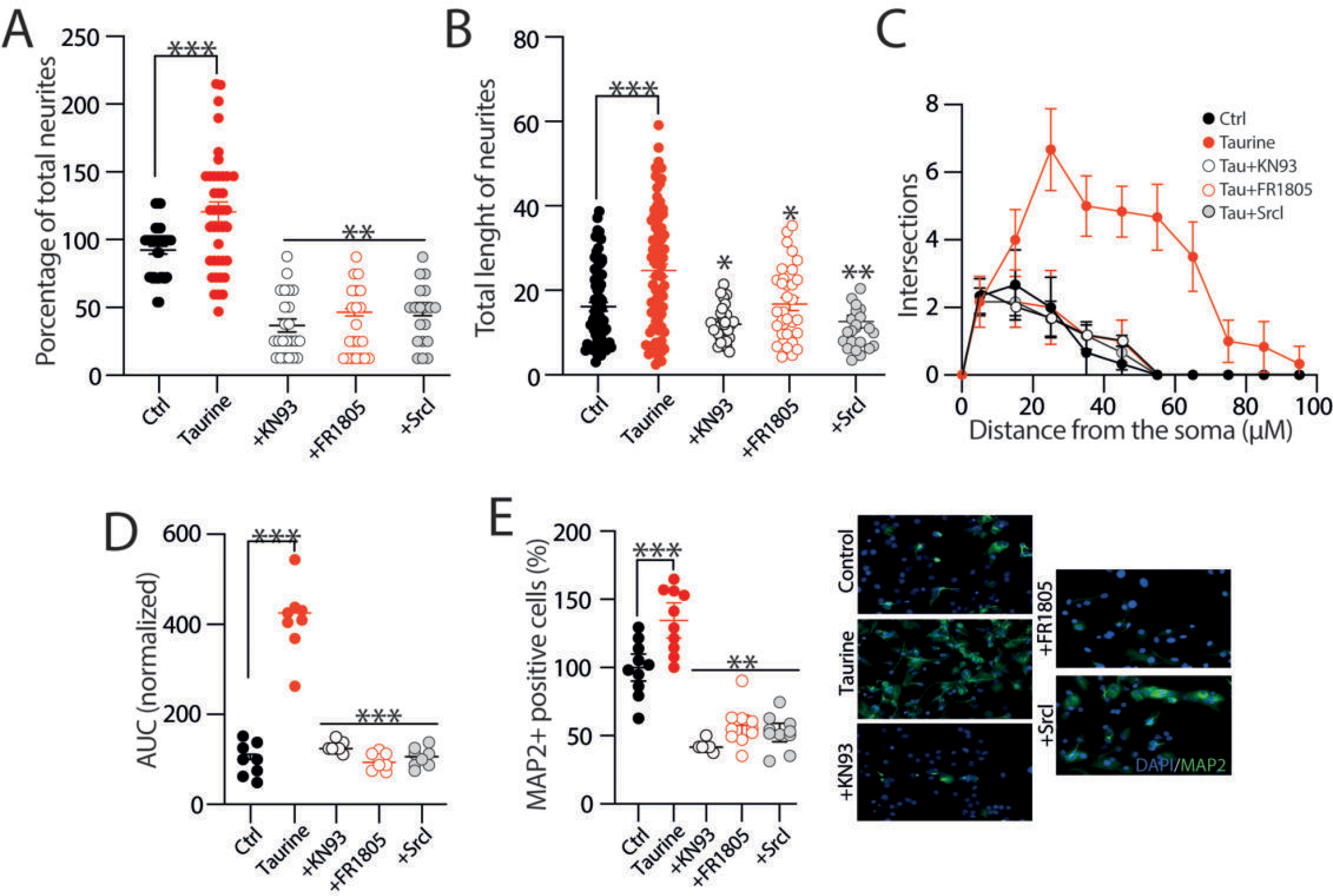
Involvement of signaling cascades in the differentiation process and neurite outgrowth of NPC SVZ mediated by taurine. **A, B)** Scatter plots showing the total number (A) and total length of neurites (B) of NPC SVZ treated with taurine or taurine + the CaMKII inhibitor KN93, taurine + the ERK1/2 kinase inhibitor FR1805, and taurine + the Src kinase inhibitor Srcl. The blockade of these signaling cascades prevented the neurite outgrowth promoted with taurine (control *n* = 25 cells; taurine *n* = 42 cells; taurine + KN93 *n* = 15 cells; taurine + FR1805 *n* = 16 cells; taurine + SrcI *n* = 19 MAP2+ cells). **C)** Sholl-like plot contrasting the number of neurite intersections in the control condition and the blockade of the indicated signaling cascades (*n* = 8 MAP2+ cells). **D)** AUC chart showing the total area obtained from the Sholl-like analysis performed in the control cells, in cells treated with taurine, and following the blockade of the indicated signaling cascades (*n* = 8 MAP2+ cells for each experimental condition). **E)** Immunofluorescence assay microphotographs against MAP2 in taurine and taurine + the signaling cascade blockers. The nuclei were stained with DAPI (blue). The blockade of CaMKII, ERK ½, or Src kinase reduced the immunoreactivity to MAP2 (*n* = 10 cell cultures for each experimental condition). \*\**p* < 0.01; \*\*\**p* < 0.001 or higher statistical significance. Error bars indicate SEM.

## Discussion

The results of this study are summarized as follows: We have provided evidence that taurine, by interacting with GABA_A_- or GABA_B_ receptors, controls the differentiation process, neurite formation, and functional expression of ionic currents of NPC SVZ. First, we demonstrated the presence of the GABA_B_ receptor subunits, GABA_B_R1 and GABA_B_R2, in NPC SVZ derived from neurospheres immunopositive to nestin and Ki-67, which are progenitor and proliferation cell markers, respectively. To the best of our knowledge, this is the first demonstration of the expression of GABA_B_R on NPC SVZ. Then, we demonstrated that taurine increases the number of DCX+ cells, favors neurite outgrowth, and promotes the development of secondary, tertiary, and higher-order neurites. These morphological changes are accompanied by the functional expression of ionic channels mediating inward and outward currents that differentially respond to the activation or blockade of GABA receptors. Since the entire maturation process was sensitive to PTX, it was mediated by the activation of GABA_A_R (see also Gutierrez-Castañeda et al., 2023). We also showed that pharmacological activation of GABA_B_R in the presence of taurine does not promote neurite outgrowth, neurite development, or functional expression of ion channels.

Contrary to this finding, the blockade of GABA_B_R with its specific antagonist CGP 35348 promoted neurite branching and stimulated the development of secondary, tertiary, and higher-order neurites of disaggregated NPC SVZ. We also demonstrated that the taurine interaction with GABA receptors involves an intracellular calcium rise that activates different signaling cascades that stimulate neurite outgrowth. Collectively, these findings show that ionotropic and metabotropic GABA receptors play bidirectional roles in the differentiation and maturation process of NPC SVZ stimulated with taurine.

### Role of GABA_A_R in the differentiation process of NPC SVZ

The role of GABA in the proliferation, differentiation, and maturation process of NPC SVZ and NPCs of the subgranular zone of the dentate gyrus has been well documented. In both neuronal niches, NPCs express the synthesizing enzyme for GABA, glutamate decarboxylase, several GABA transporters, and, in a spatiotemporal manner, GABA_A_R (Bordey et al., 2011; Liu et al., 2005; Stewart et al., 2002). In the SVZ, GABA is released in a nonsynaptic fashion, and its binding to the GABA_A_R triggers depolarization of NPCs (Liu et al., 2005; Stewart et al., 2002; Wang et al., 2003). Central to this depolarization process is the inverse chloride gradient during early neuronal development.

It is well documented that the extrusion of intracellular chloride mediated by GABA binding to the GABA_A_R leads to membrane depolarization with the subsequent activation of different voltage-sensitive channels (Agosti et al., 2017; Toth et al., 2016), including L-type calcium channels (D’Ascenzo et al., 2006; LoTurco et al., 1995; Nguyen et al., 2003). A Ca^2+^ influx via L-type channels activates calcium-dependent signaling cascades, including kinases and nuclear factors that control gene expression and cell fate. Consistent with this activity, neurons differentiated from NPCs express the Ca^2+^–calmodulin-dependent protein kinase II (CaMKII) (Kutcher et al., 2003). Activation of CaMKII phosphorylates the cellular transcription factor, the cAMP response element-binding protein (CREB) that, in turn, controls the transcription of genes critical for neurogenesis, synaptic plasticity, and neuronal survival (Lonze & Ginty, 2002; Yan et al., 2016). We have provided experimental evidence showing that the blockade of CaMKII with KN-93 inhibits neurite outgrowth and neurite branching, suggesting that calcium elevation via GABA_A_R activates CaMKII and possibly nuclear CREB.

Consistent with this possibility, CREB promotes the neuronal differentiation of NPCs (Landeira et al., 2018; Peltier et al., 2007). Likewise, the taurine-GABA_A_R interaction may activate other Ca^2+^-dependent kinases, including members of the non-receptor Src-family kinases (SFKs), a family of tyrosine kinases that regulate cell growth, differentiation, migration, and survival. Src activation phosphorylates the focal adhesion kinase (FAK) that regulates proliferation and survival via CREB phosphorylation (Schlaepfer & Hunter, 1997). The plausible participation of the Src kinases FAK and CREB in neurite outgrowth was corroborated in our study since the Src inhibitor hindered neurite development of NPC SVZ. In addition, the activation of this signaling cascade is complemented by the possible activation of PIK_3_ by Src, which, in turn, activates AKT, a kinase involved in cytoskeletal remodeling (Manning & Toker, 2017), a necessary step for dendrite development. Our experimental observations indicate that the activation of GABA_A_R with taurine triggers a depolarization that activates calcium-dependent intracellular signaling cascades that favor the development of neurites in NPCs.

### GABA_B_R in the differentiation process of SVZ NPC

Prior research has largely neglected the contribution of GABA_B_R in the differentiation process of NPCs. For example, neocortical embryonic NPCs express the GABA_B_R1 and GABA_B_R2 subunits of the GABA_B_R, and the pharmacological activation of the GABA_B_R favors the formation of neocortical-derived neurospheres (Fukui et al. 2008). Likewise, the proliferation and survival of newly born cells depend on GABA_B_R activation, and the blockade of GABA_B_R increases adult hippocampal neurogenesis (Felice et al., 2012). These observations were further corroborated by Giachino et al. (2014), who demonstrated the expression of GABA_B_R in hippocampal NPCs and that blockade of GABA_B_R increased the neuroblast proliferation and differentiation *in vivo*. These authors suggested that intracellular signaling via GABA_B_R inhibits neurogenesis and promotes the quiescence of NPCs (Giachino et al., 2014; see also Gustorff et al., 2021). Consistent with these observations, we found immunoreactivity to the GABA_B_R1 and GABA_B_R2 subunits in neurospheres derived from the SVZ. Since GABA_B_R is a metabotropic receptor, its effects should be ascribed to the dissociation of the G_i_ or G_o_ protein into G_iα_, G_oα_, and G_βγ_ subunits of the GABA_B_R complex. In such a scenario, after GABA_B_R activation, the subunit G_iα_ would inhibit the adenylyl cyclase (AC), thus decreasing the levels of cyclic adenosine 3’-5’-monophosphate (cAMP) and protein kinase A activity and decreasing the phosphorylation level of nuclear CREB (Ma et al., 2014). Likewise, the decreased levels of cAMP interfere with the functionality of the cAMP-dependent ERK/MAPK activity, and the downregulation of these kinases also negatively impacts the phosphorylation levels of CREB.

Consequently, it is reasonable to assume that the blockade of GABA_B_R with CGP55485 yields a number of positive effects related to neuronal differentiation. First, the inactivation of the G_iα_ subunit may increase the amount of cAMP and PKA activity, which would lead to increased levels of CREB phosphorylation (Lepski et al., 2013; Zhang et al., 2005). This same component may activate MAPK, another potential pathway toward CREB phosphorylation (Pearson et al., 2001). In addition, the blockade of GABA_B_R would result in transient intracellular Ca^2+^ elevation, a phenomenon that would lead to CAMKII activation and, subsequently, CREB phosphorylation (Wayman et al., 2008). These plausible mechanisms reflect the complexity and crosstalk of the signaling cascades mediated by the GABA_A_- and GABA_B_-receptors that regulate neuronal differentiation.

It is important to consider that other mechanisms may also be involved in the action of taurine in the differentiation process. For example, taurine regulates glutamate receptors and modulates intracellular Ca^2+^ signaling, which are essential for regulating neuronal growth and differentiation (Wang et al., 2015; Yang et al., 2019). These mechanisms may contribute to neuronal differentiation but require additional investigation that was beyond the scope of this study.

### Modulation of membranal currents by taurine

Another relevant finding of our study is the selective modulation of membranal currents of NPC SVZ in response to taurine treatments. Although previous studies have documented the acute effect of taurine on the membrane potential, R_N_, and action potential firing of immature and mature neurons, these effects are ascribed to an acute-chloride modulation via GABA and glycine receptors (Belluzi et al., 2004; Furukawa & Fukuda, 2023; Hosoi et al., 2022; Jiang et al., 2004; Sava et al., 2014). This possibility is unlikely in our case because NPC of the SVZ were incubated with taurine for 14 days before any experimental manipulation. One possible explanation for the electrophysiological modulation observed in our study is that taurine, in the long term, favors the adequate assembly of ionic channels rather than acutely modulating chloride conductance to alter the kinetics of the inward and outward currents. We hypothesize that this effect is an additional consequence of the differentiation capacity of taurine; however, this premise requires additional investigation.

Previous studies have demonstrated that the inward current evoked by the protocol used in our study is sensitive to TTX (Martínez-Rojas et al., 2021), which supports the notion that the taurine-mediated differentiation process of NPC SVZ involves the membranal expression of voltage-gated Na^+^ channels responsible for the action potential generation (see also Gutierrez-Castañeda, 2023). On the other hand, the kinetics of the outward currents observed in this study resemble the 4-AP-sensitive fast-inactivating K^+^ current and the TEA-sensitive delayed K^+^ current recorded in actual neurons (Griego et al., 2021, 2022). Taken together, our results suggest that taurine has the potential to selectively modulate the outward currents that determine the passive properties of excitable cells. Although additional experiments are needed to demonstrate whether taurine has the potential to stimulate the appropriate subunit assembly of ion channels, this study provides evidence that taurine has both morphogenic and electrophysiological properties to control the differentiation and maturation process of neural progenitor cells.

## Conclusion

The free amino acid taurine plays a central role in neurite outgrowth and the functional expression of ion channels that determine the passive properties of differentiated cells. Our data show that this effect occurs via GABA receptors and the subsequent activation of signaling cascades expressed by NPC SVZ. Likewise, our data suggest that GABA, acting via the ionotropic or the metabotropic receptors, has opposing effects that control adult neurogenesis in the SVZ.

**Figure 8.**
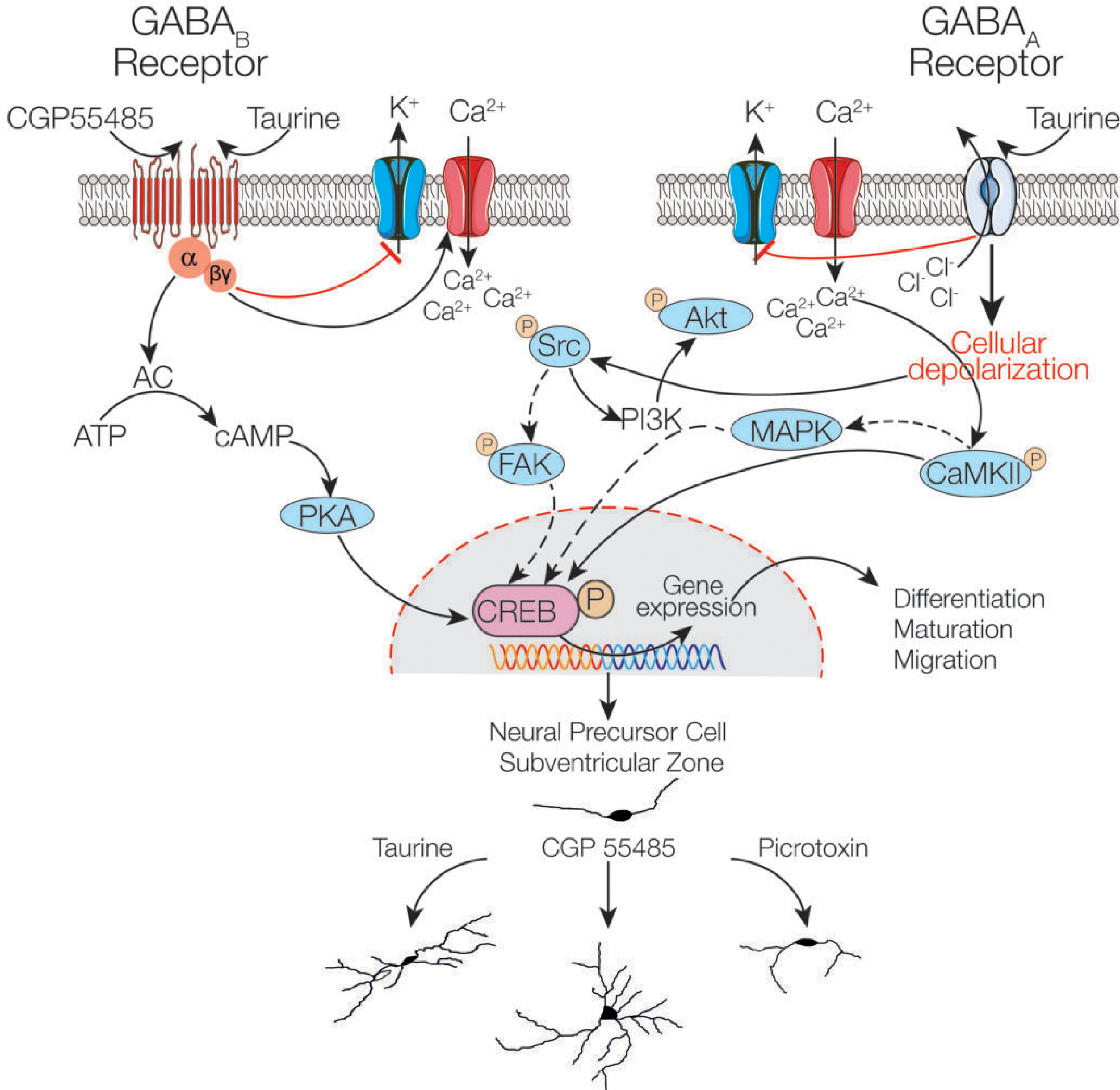
Schematic representation of the possible mechanisms necessary for the differentiation of neural precursor cells of the SVZ mediated by taurine. Taurine binds to the GABA_A_ receptor and promotes chloride extrusion and the blockade of potassium channels, favoring cellular depolarization, the activation of calcium channels, the and activation of the calcium–calmodulin (CaM)–dependent protein kinase II (CaMKII) enzyme that promotes phosphorylation of the cellular transcription factor cAMP response element-binding protein (p-CREB). Cellular depolarization also activates the non-receptor tyrosine kinase Src and phosphorylates protein kinase B or Akt, which is involved in the PI3K/AKT/mTOR signaling cascade. The blockade of the G_i_/G_o_-linked GABA_B_R with CGP55485 may increase the concentration of cAMP, possibly leading to increased CREB phosphorylation via PKA activation or MAPK cascades. Furthermore, its coupling to calcium channels through the G_βγ_ subunit of the GABA_B_R can allow high intracellular Ca^2+^ levels, activating the CAMKII signaling pathway, thus continuing the pathway mentioned with the GABAA receptors that could ultimately induce CREB activation.

## Acknowledgments

This work was supported by grants PAPIIT-UNAM IN221820 and Presupuesto Interno Facultad de Medicina (Id 49), Departamento de Investigación, Asociación para Evitar la Ceguera en México IAP Hospital “Dr. Luis Sánchez Bulnes,” and Conahcyt fellowship 783563 (NEGC).

